# The co-chaperone Fkbp5 shapes the acute stress response in the paraventricular nucleus of the hypothalamus

**DOI:** 10.1101/824664

**Authors:** Alexander S. Häusl, Jakob Hartmann, Max L. Pöhlmann, Lea M. Brix, Juan-Pablo Lopez, Elena Brivio, Clara Engelhardt, Simone Roeh, Lisa Rudolph, Rainer Stoffel, Kathrin Hafner, Hannah M. Goss, Johannes M.H.M. Reul, Jan M. Deussing, Kerry J. Ressler, Nils C. Gassen, Alon Chen, Mathias V. Schmidt

## Abstract

Disturbed activation or regulation of the stress response through the hypothalamic-pituitary-adrenal (HPA) axis is a fundamental component of multiple stress-related diseases, including psychiatric, metabolic and immune disorders. The FK506 binding protein 51 (FKBP5) is a negative regulator of the glucocorticoid receptor (GR), a main driver of HPA axis regulation, and *FKBP5* polymorphisms have been repeatedly linked to stress-related disorders in humans. However, the specific role of *Fkbp5* in the paraventricular nucleus of the hypothalamus (PVN) in shaping HPA axis (re)activity remains to be elucidated. Using deletion, overexpression, and rescue of *Fkbp5* exclusively in the PVN, we establish the fundamental importance of *Fkbp5* in the HPA axis stress response. Furthermore, we show that *Fkbp5* manipulation alters GR activation and elucidate the cellular complexity in the PVN, in which *Fkbp5* operates.

## Introduction

Life is full of challenges and appropriate coping with such events implies proper activation and termination of the stress response. The hypothalamic-pituitary-adrenal (HPA) axis is the central orchestrator of the stress response and its end product glucocorticoids (cortisol in humans, corticosterone (CORT) in rodents) mediates the adaptation to acute and chronic stressors in peripheral tissues as well as in the brain^1^. A hallmark of HPA axis regulation is the negative feedback on the secretion of stress hormones to terminate the stress response which is controlled by glucocorticoids via the GR.

A critical regulator of GR and therefore key to a successful termination of the stress response is FK506 binding protein 51 (FKBP5), which is encoded by the *FKBP5* gene^2^. FKBP5 is a Hsp90-associated co-chaperone that restricts GR function by reducing ligand binding, delaying nuclear translocation, and decreasing GR-dependent transcriptional activity ^3,4^. Hence, higher levels of *FKBP5* mRNA are associated with higher levels of circulating cortisol and reduced negative feedback inhibition of the stress response^5–9^. Consequently, GR-induced *FKBP5* levels reflect the environmental stress condition, and as such, *FKBP5* expression has been used as a stress-responsive gene marker^10^. Importantly, *FKBP5* polymorphisms have been consistently associated with stress-related psychiatric disorders such as major depression and PTSD^11–13^, where a demethylation-mediated increase in *FKBP5* expression was identified as causal in risk-allele carriers ^14^.

Despite the central importance of FKBP5 in stress system biology and stress-related disorders, detailed functional and mechanistic studies are still largely missing. Only a few human post-mortem studies focus on central tissue in order to dissect FKBP5 mechanisms, while the majority of studies use peripheral blood mononuclear cells (PBMCs) as a correlate of FKBP5 brain activity^15^. Most of the animal data were obtained from wild-type (WT) or conventional *Fkbp5* knockout mice, thereby lacking cell type-specific insights of Fkbp5 function^8,16^. To tackle this paucity of information, we here investigate the specific role of Fkbp5 in the paraventricular nucleus of the hypothalamus (PVN) the key brain region orchestrating the stress response. Using site-specific manipulations of *Fkbp5*, single-cell RNA expression profiling, and functional downstream pathway analyses, our data unravel a key role of PVN Fkbp5 in shaping the body’s stress system (re)activity, with important implications for its contribution to stress-related disorders.

## Results

### Loss of *Fkbp5* in the PVN alters HPA axis physiology

To study the effects of Fkbp5 in the PVN, we first generated PVN-specific conditional *Fkbp5* knockout mice (*Fkbp5*^PVN−/−^) by crossing *Fkbp5*^lox/lox^ mice (generated in-house; for details see online methods) with the Sim1-Cre mouse line, which expresses *Cre* recombinase in Sim1^+^ neurons mostly concentrated within the PVN (Figure 1A). The successful deletion of *Fkbp5* in the PVN was assessed by mRNA and protein analysis (Figure 1B, Supplementary Figure 1). Under basal conditions, adult mice lacking Fkbp5 in the PVN showed significantly lower body-, and adrenal weights, and higher thymus weights compared to their WT littermates (Figure 1C). Interestingly, *Fkbp5*^PVN−/−^ mice in young adolescence showed no significant differences in body weight,adrenal or thymus weights (Supplementary Figure 2), indicating an age-dependent phenotype of Fkbp5 in the HPA-axis’ response.

**Figure 1:**
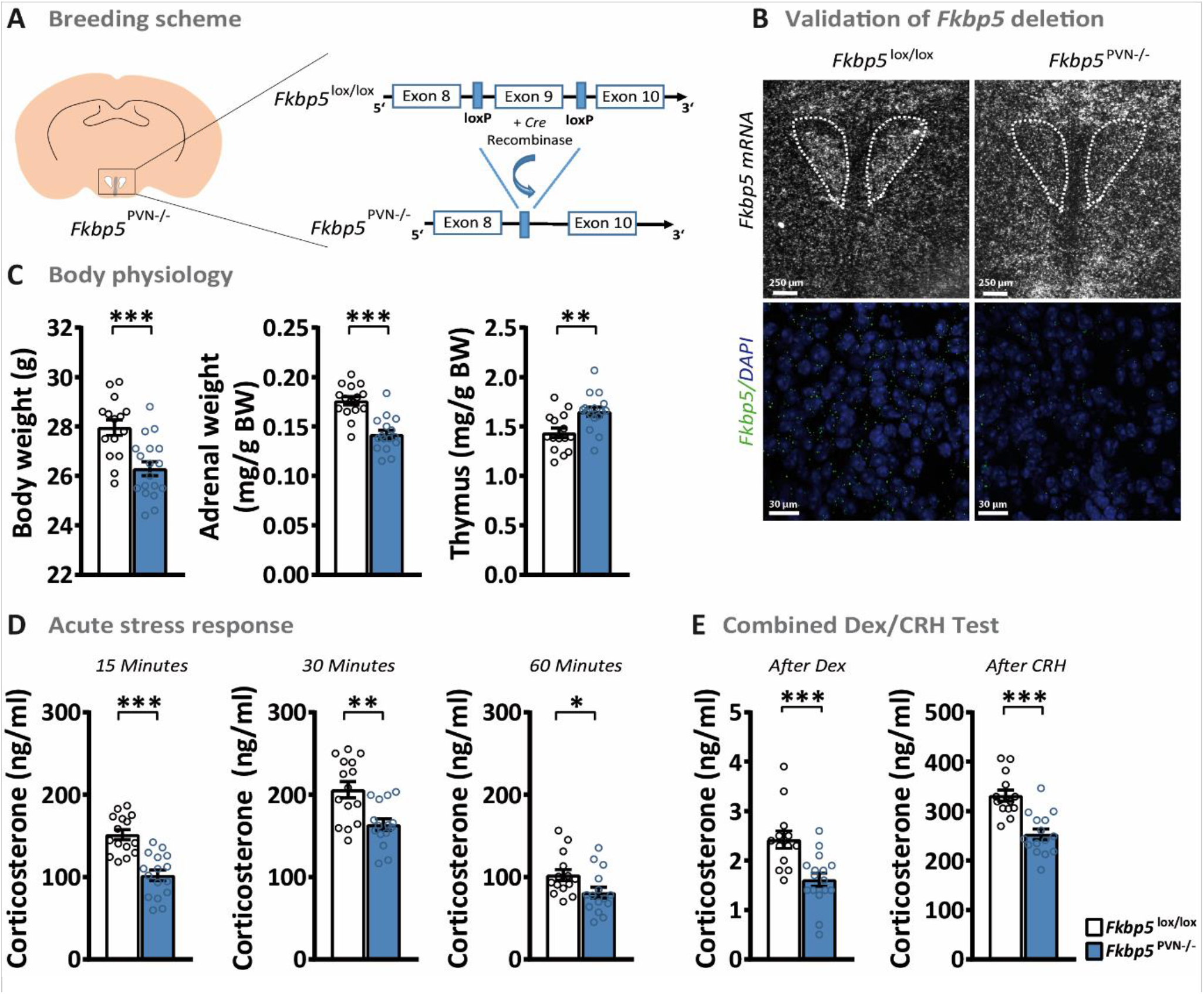
Loss of *Fkbp5* in the PVN alters HPA axis physiology. **(A)** Cre-LoxP based generation of the *Fkbp5*^PVN−/−^ mouse line. **(B)** Validation of *Fkbp5* mRNA expression in the PVN via *In-Situ* hybridization (ISH) and RNAscope. **(C)** *Fkbp5*^PVN−/−^ mice (n = 16) presented reduced body weight, lowered adrenal weights and increased thymus weights under non-stressed conditions compared to their WT littermates (n = 15). **(D)** Corticosterone levels were significantly reduced following a 15 minutes restraint stress until at least 60 minutes after stress onset. **(E)** A combined Dex/CRH test showed a significantly pronounced response to a low dose of dexamethasone as well as a dampened response to CRH injection. Data are presented as mean ± SEM. All data were analyzed with a student’s t-test. * = p < 0.05, ** = p < 0.01 and *** = p < 0.001.

As PVN *Fkbp5* mRNA levels are highly responsive to an acute stressor^10^, we hypothesized that *Fkbp5*^PVN−/−^ mice have an altered stress response following an acute challenge. Under basal conditions during the circadian trough, no differences in corticosterone secretion were detected in young and adult mice (Supplementary Figure 1 and 2). However, already after 15 minutes of restraint stress *Fkbp5*^PVN−/−^ displayed significantly reduced plasma corticosterone levels compared to the control group (Figure 1D). The dampened stress response was persistent for 60 minutes after stress onset, while levels of the adrenocorticotropic hormone (ACTH) were not altered under basal or acute stressed conditions (Supplementary Figure 1).

To further investigate the effect of Fkbp5 on GR sensitivity, we performed a combined dexamethasone (Dex, a synthetic GC) – corticotropin-releasing hormone (CRH) test. The combined Dex/CRH test is a method to analyze HPA axis (dys)function in depressed individuals or animals, measuring the responsiveness of the body’s stress response system through suppression (by Dex injection) and stimulation (by CRH injection) of the HPA axis^17^. The injection of a low dose of Dex (0.05 mg/kg) resulted in a reduction in blood corticosterone levels compared to the evening corticosterone levels in both groups. Interestingly, mice lacking *Fkbp5* in the PVN showed 1.5-fold lower levels of corticosterone compared to their WT littermates. Following CRH stimulation (0.15 mg/kg) *Fkbp5*^PVN−/−^ mice showed a significantly lower reaction to CRH than control mice (Figure 1E). These data indicate that specific deletion of *Fkbp5* in the PVN dampens HPA axis response and enhances GR sensitivity.

### Overexpression of *Fkbp5* in the PVN induces a stress-like phenotype in C57Bl/6 mice under basal conditions

Chronic or acute stress upregulates *Fkbp5* in distinct brain regions, such as the PVN^10^. Therefore, we were interested to explore whether selective overexpression of *Fkbp5* in the PVN would be sufficient to affect body physiology and stress system regulation. To do so, we bilaterally injected 200 nl of an adeno-associated virus (AAV) containing a *Fkbp5* overexpression vector into the PVN of young adult (10 weeks) C57Bl/6 mice (*Fkbp5*^PVN OE^, Figure 2A-B). The AAV-mediated overexpression resulted in a (4-fold) increase in *Fkbp5* mRNA and protein levels (Fkbp5) in the PVN (Supplementary Figure 3).

**Figure 2:**
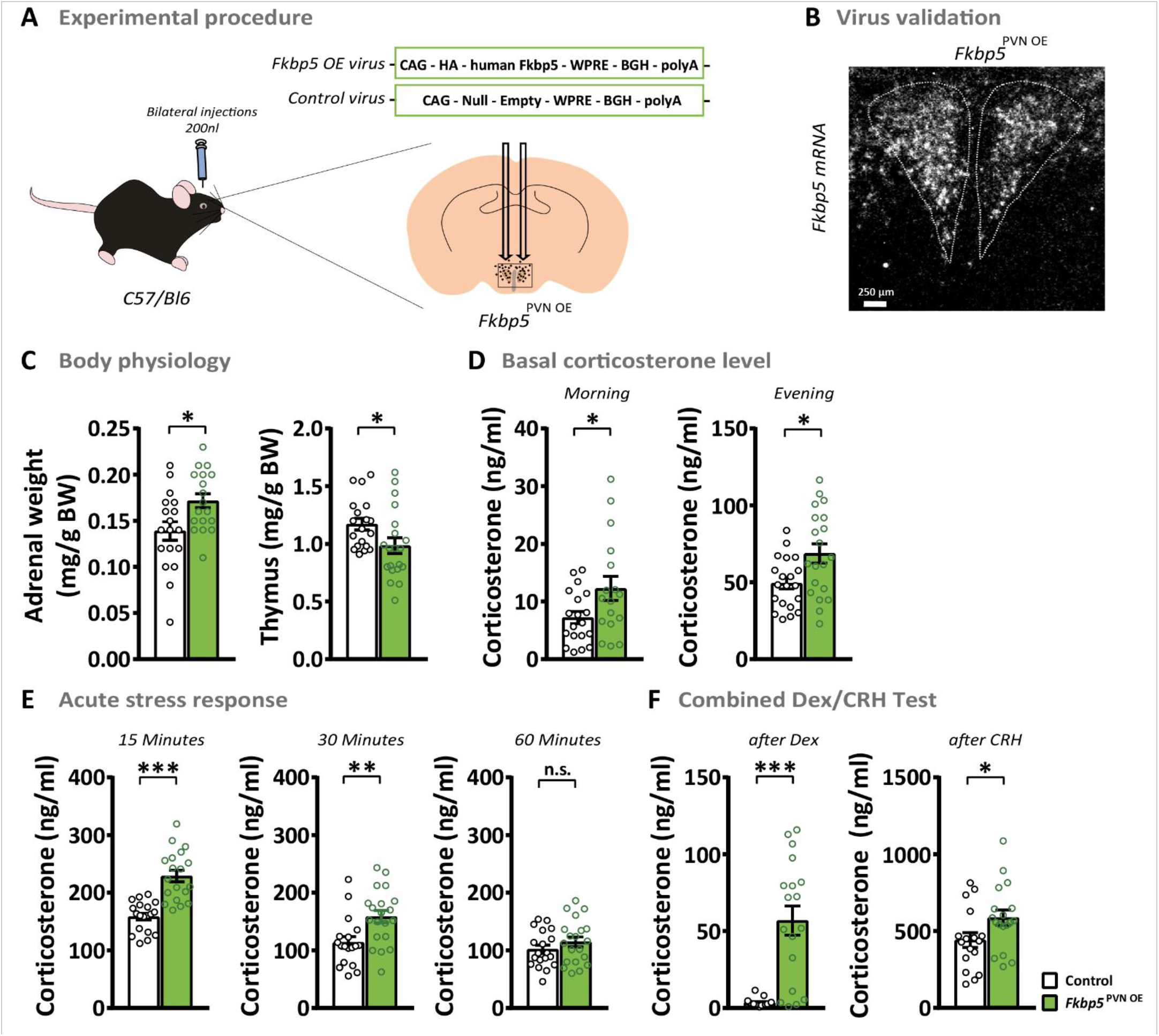
The overexpression of *Fkbp5* in C57Bl/6 mice induces stress-like phenotype under basal conditions. **(A)** Overexpression of *Fkbp5* in the PVN was achieved by bilateral viral injections. **(B)** Validation of *Fkbp5* mRNA overexpression in the PVN by ISH. **(C)** *Fkbp5*^PVN^ OE mice (n = 20) showed significantly increased adrenal weights and a reduced thymus weight under non-stressed conditions compared to the controls (n = 20). **(D)** *Fkbp5* overexpression resulted in heightened corticosterone levels during the day. **(E)** 15 and 30 minutes after stress onset, *Fkbp5*^PVN OE^ mice displayed significantly higher corticosterone levels. **(F)** *Fkbp5*^PVN OE^ mice showed significantly elevated corticosterone 6 h after dexamethasone treatment. The following CRH injection further significantly increased the corticosterone release compared to controls. Data are presented as mean ± SEM. All data were analyzed with student’s t-test. * = p < 0.05, ** = p < 0.01, *** = p < 0.001, n.s. = not significant.

Intriguingly, *Fkbp5* overexpression altered the physiology of stress responsive organs. *Fkbp5*^PVN OE^ animals showed a significantly reduced thymus weight and increased adrenal weights compared to their littermates (Figure 2C), the hallmark of chronically stressed animals^18,19^. Furthermore, overexpression of *Fkbp5* affected the circadian rhythm of corticosterone secretion, indicated by increased blood corticosterone levels in the morning as well as the evening (Figure 2D). Consequently, ACTH levels of *Fkbp5*^PVN OE^ animals were also increased under non-stressed conditions (Supplementary Figure 3). Next, we analyzed distinguished stress markers under basal conditions in order to determine whether consequences of PVN-specific *Fkbp5* overexpression are also detectable at the molecular level. Interestingly, *Nr1c3* and *Crh* mRNA expression in the PVN were increased in *Fkbp5*^PVN OE^ animals compared to controls, whereas *Avp* mRNA levels were not altered (Supplementary Figure 3). Together, these results are comparable to chronically stressed animals and demonstrate that local overexpression of *Fkbp5* in the PVN is sufficient to mimic a stress-like phenotype without physically challenging the animals.

In accordance to the knock-out studies with the *Fkbp5*^PVN−/−^ animals, we investigated the endocrinology of *Fkbp5*^PVN OE^ animals after an acute challenge. As expected, we detected higher blood corticosterone levels in *Fkbp5*^PVN OE^ mice 15 and 30 minutes after stress onset compared to the stressed controls (Figure 2E). However, we could not detect any difference in ACTH release (Supplementary Figure 3). No differences between both groups were observed at 60 and 90 minutes after restraint stress (Figure 2E, Supplementary Figure 3). These data show that *Fkbp5*^PVN OE^ mice have a hyperactive HPA axis response and are more vulnerable to acute stress exposure.

To further assess GR sensitivity in *Fkbp5* overexpressing animals, we again tested the response to a combined Dex/CRH test. While control animals showed a decline (< 5 ng/ml) in blood corticosterone levels 6 h after Dex injection, *Fkbp5*^PVN OE^ mice showed almost no response to Dex treatment (Figure 2F). Interestingly, the subsequent CRH injection resulted in a higher corticosterone release in *Fkbp5*^PVN OE^ mice compared to controls (Figure 2F). These results suggest that excess levels of Fkbp5 in the PVN lead to a decreased GR sensitivity and thereby to an altered HPA axis response. Taken together, animals overexpressing Fkbp5 in the PVN show a hyperactive function of the HPA axis under basal and acute stress conditions, thereby mimicking the physiological hallmarks of chronic stress exposure and HPA axis hyperactivity, as observed in multiple stress-related diseases^20^.

### Reinstatement of endogenous *Fkbp5* in the PVN of global *Fkbp5* knock-out animals normalizes the body’s stress response

Global loss of *Fkbp5* results in a more sensitive GR and better coping behavior of mice after stress^8,15,18,21^. Our results showed that *Fkbp5* in the PVN is necessary for an undisturbed stress system function. To investigate whether *Fkbp5* function in the PVN is sufficient to restore a “normal” HPA axis (re)activity, we aimed to re-instate native *Fkbp5* expression in global *Fkbp5* knock-out animals selectively in the PVN. Therefore, we injected a Flp recombinase expressing virus into *Fkbp5*^Frt/Frt^ mice. These mice carry a FRT flanked reporter selection (stop) cassette within the *Fkbp5* locus, leading to a disruption of Fkbp5. Flp removes the stop cassette from the *Fkbp5* locus, resulting in endogenous Fkbp5 re-expression (Figure 3A-B, Supplementary Figure 4). In parallel to the two previous mouse models, we assessed body physiology, basal corticosterone levels, and the acute stress response. Interestingly, mice with re-instated *Fkbp5* expression (*Fkbp5*^PVN Rescue^) showed significantly higher adrenal weights as compared to their control littermates (Figure 3C). Furthermore, we observed that the reinstatement of *Fkbp5* in the PVN resulted in significantly increased blood CORT levels in the morning under basal conditions (Figure 3D), with no effect on thymus weights, evening CORT, and basal ACTH levels (Supplementary Figure 4). ISH analysis revealed significant higher levels of *Crh* mRNA, but no changes in *Nr1c3* and *Avp* mRNA expression in the PVN (Supplementary Figure 4). Next, we monitored blood corticosterone levels after 15 minutes of restraint stress. Here, we observed significantly higher CORT levels 15 and 60 minutes after stress onset (Figure 3E). Intriguingly, no differences were detected in the combined Dex/CRH test (Supplementary Figure 4), indicating that a PVN-driven over-activation of the HPA axis is necessary for a desensitization of GRs in the PVN and the pituitary. These rescue experiments underline the importance of Fkbp5 in the acute stress response and demonstrate that Fkbp5 in the PVN is necessary and sufficient to regulate HPA axis (re)activity.

**Figure 3:**
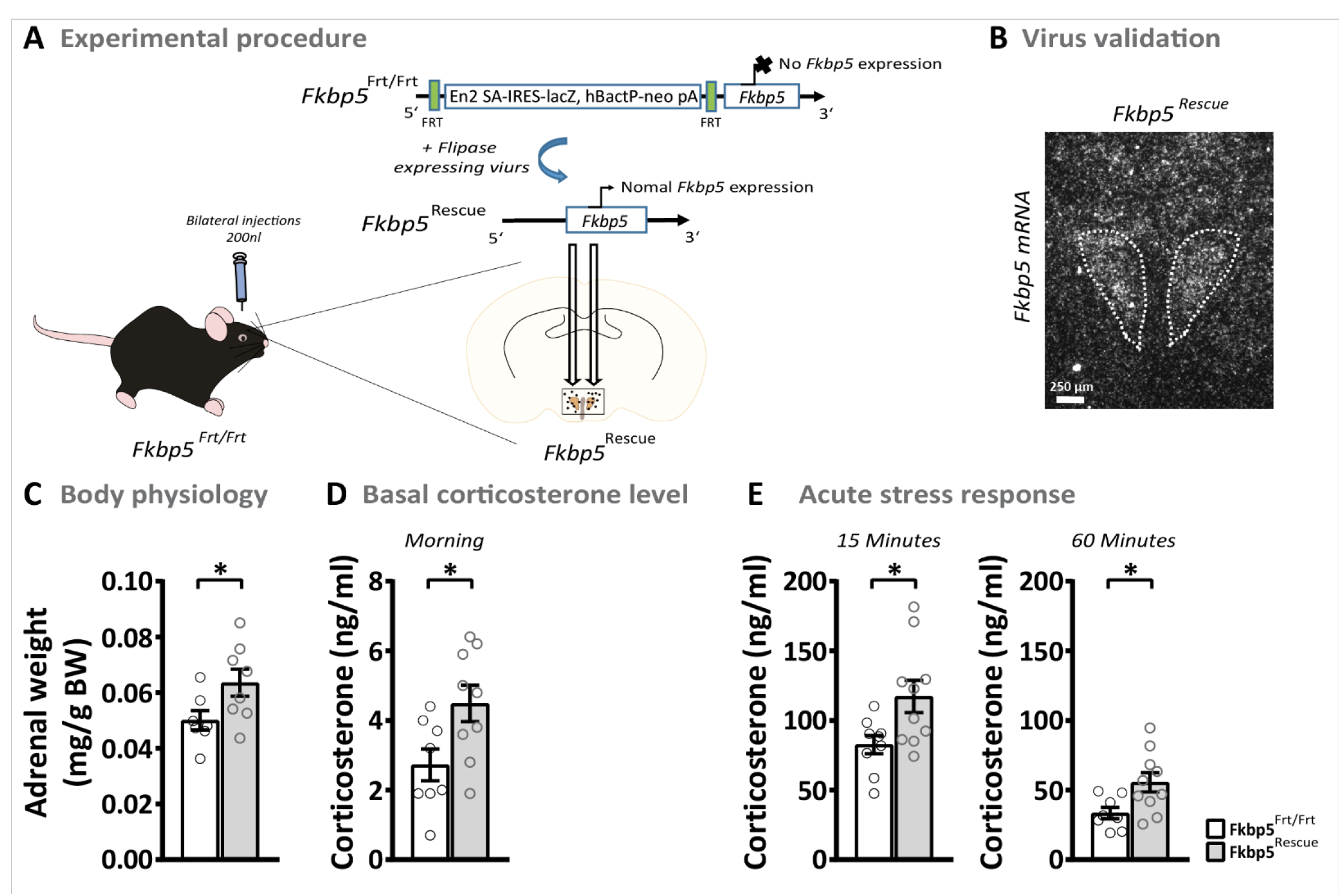
The reinstatement of endogenous *Fkbp5* in the PVN of global Fkbp5 knock-out animals. **(A)** Experimental procedure. **(B)** Validation of successful *Fkbp5* rescue by ISH. **(C)** The reinstatement of *Fkbp5* in the PVN resulted in increased adrenal weights and elevated morning corticosterone levels under basal conditions **(D)**. Furthermore, *Fkbp5* re-instated animals displayed a significantly higher corticosterone response after restraint stress **(E)**. Data are presented as mean ± SEM. All data were analyzed with student’s t-test. * = p < 0.05.

### *Fkbp5* manipulation directly affects GR phosphorylation

It is well known that ligand-binding induced phosphorylation of GR plays an important role in response to hormone signaling^22^. The main phosphorylation sites involved in hormone signaling of GR are at Serine(Ser)^203^ (mouse S^212^), Ser^211^ (mouse Ser^220^) and Ser^226^ (mouse Ser^234^) and are associated with GR activity^22,23^. Here, we tested the hypothesis that the co-chaperone Fkbp5 regulates phosphorylation of GR in *Fkbp5*^PVN−/−^ and *Fkbp5*^PVN OE^ mouse lines. To do so, we dissected the PVN of *Fkbp5*^PVN−/−^ and *Fkbp5*^PVN OE^ mice and measured the phosphorylation levels of Ser^203^, Ser^211^, and Ser^234^ under basal and stress conditions.

Under basal conditions, animals lacking Fkbp5 in the PVN showed significantly less GR phosphorylation at Ser^203^ (Figure 4A). Furthermore, *Fkbp5*^PVN−/−^ animals displayed higher phosphorylation of GR at Ser^234^ and Ser^211^ in comparison to their WT littermates (Figure 4B-C). Under stressed conditions, deletion of Fkbp5 had the same effects on pGR^Ser211^ and pGR^Ser234^ as we observed under basal conditions (Figure 4B-C). Levels of pGR^Ser203^ were found to be unchanged in the *Fkbp5*^PVN−/−^ after an acute stress compared to the basal levels. However, pGR^Ser203^ levels of the control group increased after stress (Figure 4A).

**Figure 4:**
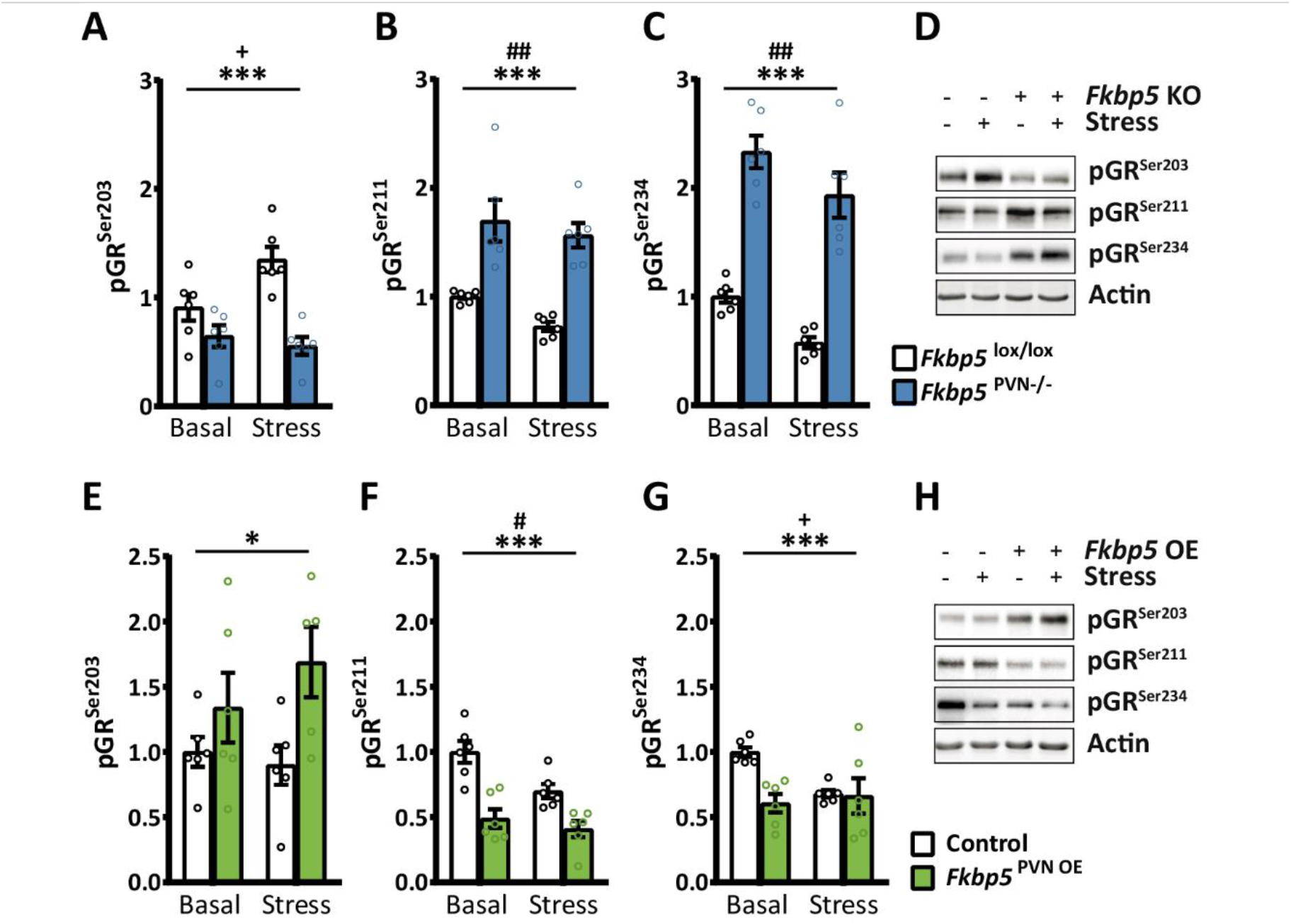
Fkbp5 manipulation affects phosphorylation of the glucocorticoid receptor (GR). **(A)** Animals lacking Fkbp5 in the PVN showed significantly lower phosphorylation at pGR^Ser203^ and higher levels of pGR^Ser211^ **(B)** and pGR^Ser234^ **(C)** compared to the control animals. Fkbp5^PVN OE^ animals showed the opposite effect on GR phosphorylation with higher phosphorylation on pGr^Ser203^ **(E)**. Additionally, we observed a significantly lower phosphorylation at the GR sites Ser^211^ **(F)** and Ser^234^ **(G)**. Representative blots are shown in **(D)** and **(H)**. Group size for A-H: 6 vs. 6. Data are presented as mean ± SEM and were analyzed with a two-way ANOVA. * = significant genotype effect, (* = p < 0.05, ** = p < 0.01, *** = p < 0.001). + = significant genotype x stress interaction (+ = p < 0.05),(# = significant stress effect (# = p < 0.05, ## = p < 0.01).

Intriguingly, *Fkbp5*^PVN OE^ animals showed exactly the opposing phenotype at all three phosphorylation sites with less pGr^Ser234^ and pGr^Ser211^ and higher phosphorylation at Ser^203^ under basal conditions (Figure 4E-G). In parallel to the unstressed condition, the overexpression of *Fkbp5* resulted in less GR phosphorylation at Ser^211^ and higher levels of pGR^Ser203^ compared to their control group after stress (Figure 4E-F). Interestingly, levels of pGR^Ser234^ were unchanged after stress (Figure 4G). Notably, total GR levels were not significantly altered in both experimental groups and conditions (Supplementary Figure 5). Despite the altered GR phosphorylation, we could not detect any significant changes in GR enrichment at the glucocorticoid response element (GRE) in the *Crh* gene after acute stress (Supplementary Figure 6), which may be due to the use of an antibody that recognizes all GR molecules irrespective of its phosphorylation state.

Overall our data demonstrate that *Fkbp5* manipulation in the PVN affects GR phosphorylation at all three major phosphorylation sites. Furthermore, it suggests that Fkbp5 affects GR phosphorylation and that the changed phosphorylation, at least partly, leads to the corticosterone phenotype in *Fkbp5*^PVN−/−^ and *Fkbp5*^PVN OE^ animals by altering GR activity.

### Fkbp5 in the PVN acts in a complex cellular context

To further unravel the expression profile of Fkbp5 in the PVN and to detect cellular populations that might be mediating the effects of Fkbp5 on HPA axis control, we used a single-cell RNA sequencing dataset consisting of 5,113 single cells isolated from the PVN. The single-cell expression data reveal a complex cellular composition, with the majority of cells identified as neurons (38%), ependymal cells (25%) and astrocytes (14%) (Figure 5A-B). *Fkbp5* was found to be differentially and cell-type specifically expressed, with the biggest *Fkbp5*+ cell population found in GABAergic neurons (42%). A significant expression of *Fkbp5* was also detected in neuronal populations known to be directly involved in HPA axis regulation, most prominently in *Crh* positive neurons (Figure 5C). However, it is important to point out that the expression levels of *Fkbp5* are relatively low. Unfortunately, lowly expressed genes may not be detected using this technique^24^ and therefore many *Fkbp5* positive cells may have remained undetected in this dataset. To circumvent this problem, we next performed a targeted co-expression study of *Fkbp5* with five major markers that are characteristic of the stress response oxytocin (*Oxt*), somatostatin (*Sst*), vasopressin (*Avp*), thyrotropin releasing hormone (*Trh*), and *Crh* under basal and stress conditions (Figure 6). We observed a strong but not complete co-localization of *Fkbp5* expression with these neuropeptide-expressing cellular populations in the PVN under basal conditions. Interestingly, a detailed quantification of the change in co-expression following stress revealed that there was a significant increase in *Crh-Fkbp5* co-localization only in the *Crh*-expressing neurons (Figure 6, Suppl. Fig 7).

**Figure 5:**
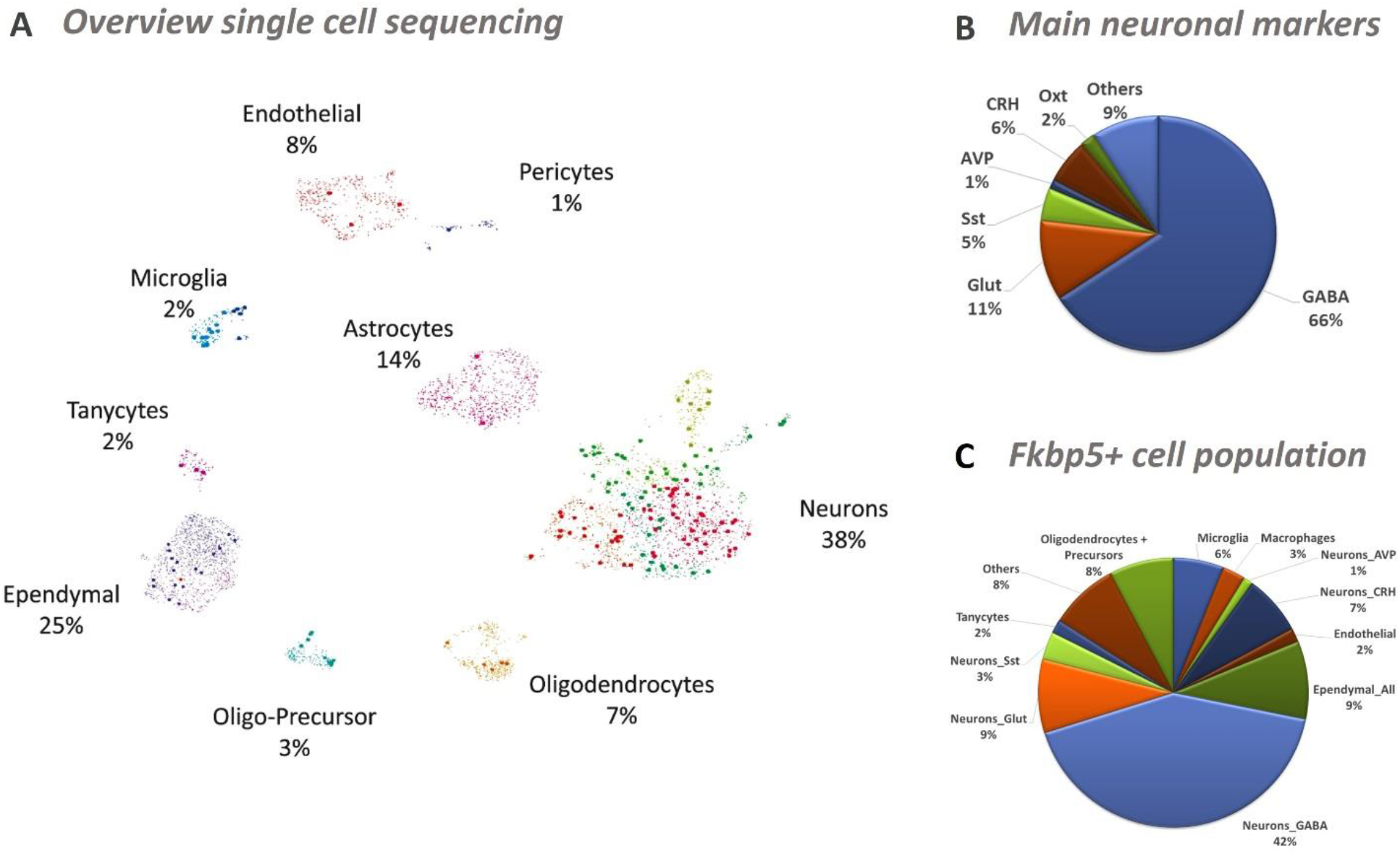
Single-cell RNA sequencing of cells in the PVN under non-stressed conditions. **(A)** Single-cell sequencing depicted several different cell types. With the majority being neurons (38%), ependymal cells (25%) and astrocytes (14%). Fkbp5^+^ cells are highlighted. **(B)** Neurons could be divided mostly into GABAergic (66%) and glutamatergic (Glut, 11%) cells. Furthermore, the well-known stress markers corticotropin-releasing hormone (*Crh*, 6%), somatostatin (*Sst*, 5%), oxytocin (*Oxt*, 2%) and vasopressin (*Avp*, 1%) could be detected under basal conditions. **(C)** Diversity of Fkbp5^+^ cell population.

**Figure 6:**
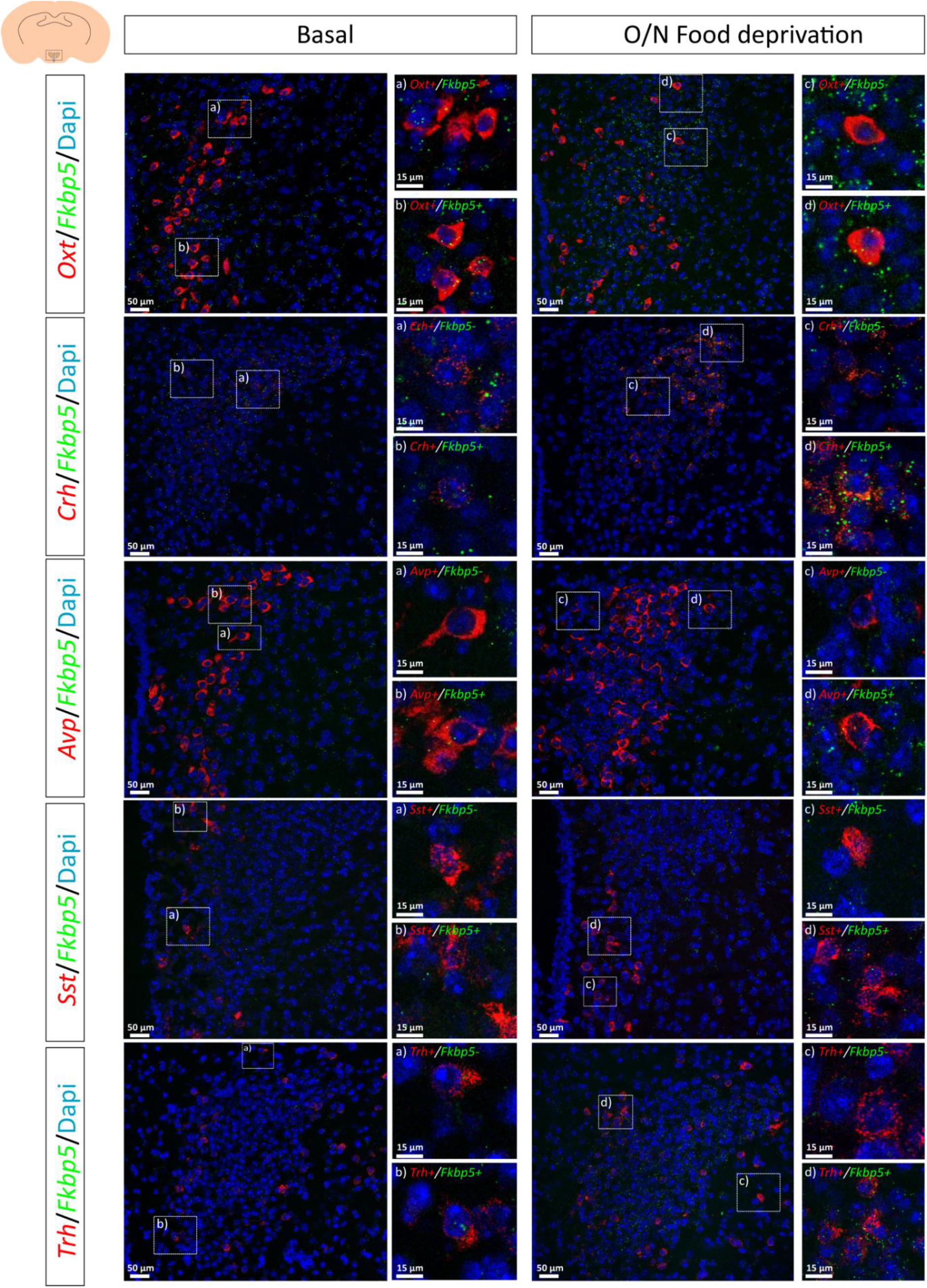
RNAscope analysis of *Fkbp5* expression in stress response neuronal cell populations under basal and stressed conditions. In comparison to single-cell sequencing, RNAscope revealed a significant higher co-localization ratio of *Fkbp5* in oxytocin (*Oxt*), corticotropin-releasing hormone (*Crh*), vasopressin (*Avp*), somatostatin (*Sst*) and thyronine releasing hormone (*Trh*) neurons. Overnight food deprivation increased *Fkbp5* mRNA in all cell populations.

These data reveal the complex cell type specific expression pattern of *Fkbp5* under stress and basal conditions in the PVN. Furthermore, they indicate a crucial role of *Fkbp5* in *Crh* positive neurons after an acute stress challenge.

## Discussion

FKBP5 was first associated with stress-related disorders in 2004^13^ and has been studied extensively over the past 15 years with regard to stress regulation and sensitivity. However, detailed cell-type and region-specific manipulations of Fkbp5 in the brain are still lacking. In this study, we highlight the importance of this co-chaperone in the regulation of the acute stress response through the combined analysis of deletion, overexpression, and rescue of *Fkbp5* exclusively in the PVN.

*Fkbp5* is a stress responsive gene and past research has shown that its main effects occur after chronic or acute stress^8,16,21,25,26^. Given that deletion of *Fkbp5* in the PVN mimics the previously described phenotype of *Fkbp5* KO mice^21^, with regard to their basal neuroendocrine profile and HPA axis function, our data illustrate that the functional contribution of Fkbp5 to HPA axis activity is centered in the PVN. In addition, reinstatement of native basal *Fkbp5* expression in the PVN of *Fkbp5* KO mice was sufficient to normalize HPA axis function. Interestingly, the phenotype of intensified HPA axis suppression due to the loss of *Fkbp5* in the PVN emerges only in adult animals, excluding developmental effects and underlining the previously reported importance of Fkbp5 in ageing^27,28^.

Further, our results highlight the essential role of Fkbp5 in stress adaptation, as PVN-specific Fkbp5 excess is sufficient to reproduce all physiological and endocrinological hallmarks of a chronic stress situation ^29,30^. Interestingly, the results of our *Fkbp5*^PVN OE^ cohort are comparable to the neuroendocrine effect of GR deletion in the PVN^31^. The consequence of a heightened *Fkbp5* expression in the PVN is two-fold. Firstly, it leads to direct changes in GR sensitivity and downstream GR signaling directly in the PVN. Secondly, PVN *Fkbp5* overexpression dramatically affects GR sensitivity and feedback at the level of the pituitary, as demonstrated by the inability of a low Dex dose (that does not cross the blood-brain barrier) to suppress corticosterone secretion. This secondary effect is likely due to the constant overproduction of CRH in the PVN and very similar to the effects of HPA hyperactivity observed in many depressed patients^17,32^.

Mechanistically, we explored the role of Fkbp5 in modulating GR phosphorylation. The status of GR phosphorylation at Ser^211^, Ser^203^, and Ser^234^ is associated with transcriptional activity, nuclear localization, and ability to associate with GRE containing promoters^22^. Whereas higher levels of phosphorylation at Ser^211^ are associated with full transcriptional activity and localization in the nucleus, increased phosphorylation of Ser^203^ is linked to a transcriptional inactive form of GR within the cytoplasm and thereby less active GR^33–35^. In our experiments, overexpression of Fkbp5 resulted in dephosphorylation at Ser^211^ and higher phosphorylation at Ser^203^, suggesting that the GR is mostly located in the cytoplasm and less active. Deletion of Fkbp5 showed the opposing effect, indicating a more active GR in *Fkbp5*^PVN−/−^ animals. It has previously been reported that GR phosphorylation is regulated by several kinases, including CDK5 and ERK^22,36^, and Fkbp5 has also been shown to be associated with CDK5 in the brain^37^. Therefore, we hypothesize that Fkbp5 also interacts with CDK5 to phosphorylate GR at multiple phosphorylation sites, thereby directly affecting ligand-dependent GR activity.

Given the complexity of the different cell types with highly specialized functions in the brain, it is essential to gain a deeper understanding of the cellular architecture of the PVN and the specific function of Fkbp5 in this context. Previously, it was assumed that Fkbp5 is quite widely expressed in most cell types of the nervous system^38^. However, our current data suggest that while *Fkbp5* is indeed expressed in the PVN, it is enriched in specific sub-populations, including for example GABAergic neurons, *Crh*^+^ neurons and microglia, but largely absent in others, such as astrocytes and endothelial cells. Interestingly, when quantifying Fkbp5 regulation, we identified a highly selective regulation of *Fkbp5* in *Crh*^+^ neurons, further supporting the central role of Fkbp5-controlled GR feedback in this neuronal subpopulation.

In summary, this study is the first to specifically manipulate Fkbp5 in the PVN and underlines its central importance in shaping HPA axis regulation and the acute stress response. The results have far-reaching implications for our understanding of stress physiology and stress-related disorders.

## Methods

### Animals & animal housing

All experiments were performed in accordance with the European Communities’ Council Directive 2010/63/EU. The protocols were approved by the committee for the Care and Use of Laboratory animals of the Government of Upper Bavaria, Germany. The mouse lines *Fkbp5*^lox/lox^ and *Fkbp5*^PVN−/−^ and *Fkbp5*^Frt/Frt^ were generated in house. Male mice aged between 3-5 months were used for all experiments. During the experimental time, animals were kept singly housed in individually ventilated cages (IVC; 30cm × 16cm × 16cm; 501cm^2^), serviced by a central airflow system (Tecniplast, IVC Green Line – GM500). Animals were maintained on a 12:12hr light/dark cycle, with constant temperature (23±2°C) and humidity of 55% during all times. Experimental animals received *ad libitum* access to water and standard research diet (Altromin 1318, Altromin GmbH, Germany) and IVCs had sufficient bedding and nesting material as well as a wooden tunnel for environmental enrichment.

### Generation of *Fkbp5* mice

Conditional *Fkbp5* knockout mice are derived from embryonic stem cell clone EPD0741_3_H03 which was targeted by the knockout mouse project (KOMP). Frozen sperm obtained from the KOMP repository at UC Davis was used to generate knockout mice (*Fkbp5^tm1a(KOMP)Wtsi^*) by *in vitro* fertilization. These mice designated as *Fkbp5*^Frt/Frt^ are capable to re-express functional Fkbp5 upon Flp recombinase-mediated excision of a frt-flanked reporter-selection cassette integrated in the *Fkbp5* gene. Mice with a floxed *Fkbp5* gene designated as *Fkbp5*^lox/lox^ (*Fkbp5^tm1c(KOMP)Wtsi^*) were obtained by breeding *Fkbp5*^Frt/Frt^ mice to Deleter-Flpe mice^39^. Finally, mice lacking *Fkbp5* in PVN neurons (*Fkbp5*^PVN−/−^) were obtained by breeding *Fkbp5*^lox/lox^ mice to Sim1-Cre mice^40^. Genotyping details are available upon request.

### Viral overexpression and rescue of *Fkbp5*

For overexpression and rescue experiments, stereotactic injections were performed as described previously^41^. We used an adeno-associated bicistronic AAV1/2 vector for the overexpression and rescue studies. In the overexpression experiments, the vector contained a CAG-HA-tagged-FKBP51-WPRE-BGH-polyA expression cassette (containing the coding sequence of human Fkbp51 NCBI CCDS ID CCDS4808.1). The same vector construct without expression of Fkbp5 (CAG-Null/Empty-WPRE-BGH-polyA) was used as a control. Virus production, amplification, and purification were performed by GeneDetect. For the rescue experiment, a viral vector containing a flippase expressing cassette (AAV2-eSYN-eGFP-T2A-FLPo, Vector Biolabs; VB1093) was used to induce endogenous *Fkbp5* expression in *Fkbp5*^FRT/FRT^ mice. Control animals were injected with a control virus (AAV2-eSYN-eGFP; Vector Biolabs; VB1107). For both experiments, mice were anesthetized with isoflurane, and 0.2 μl of the above-mentioned viruses (titers: 1.6 × 10^12–13^ genomic particles/ml) were bilaterally injected in the PVN at 0.05 μl/min by glass capillaries with tip resistance of 2–4MΩ in a stereotactic apparatus. The following coordinates were used: −0.45 mm anterior to bregma, 0.3 mm lateral from midline, and 4.8 mm below the surface of the skull, targeting the PVN. After surgery, mice were treated for 3 d with Metacam via i.p. injections and were housed without any experiments for 3-4 weeks for total recovery. Successful overexpression and reinstatement of *Fkbp5* was verified by *ISH*.

### Acute stress paradigm

For acute stress, mice were placed in a 50 ml falcon tube with 2 holes at the top and the lid to provide enough oxygen and space for tail movement. On the experimental day, each animal was placed in the 50 ml falcon for 15 minutes in their individual home cage. After 15 minutes, animals were removed from the tube and the first blood sample was collected by tail cut. Until the following tail cuts at 30, 60 and 90 minutes after stress onset, the animals remained in their home cage to recover. Basal CORT levels (morning CORT) were collected one week prior the acute stress paradigm at 8 am.

### Combined Dex/CRH test

To investigate HPA axis function we performed a combined Dex/CRH test as described previously^8^. On the experimental day, mice were injected with a low dose dexamethasone (0.05 mg/kg, Dex-Ratiopharm, 7633932) via i.p. injections at 9 am in the morning. 6 hours after Dex injection, a blood sample was collected via tail cut (after Dex value), followed by an injection of CRH (0.15 mg/kg, CRH Ferrin Amp). 30 min after CRH injection, another blood sample was obtained (after CRH value). All samples from the acute stress experiments and the Dex/CRH test were collected in 1.5 ml EDTA-coated microcentrifuge tubes (Sarstedt, Germany). All blood samples were kept on ice and centrifuged for 15 min at 8000 rpm and 4°C. Plasma samples were transferred to a new, labeled microcentrifuge tubes and stored at −20 °C until further processing.

### Sampling procedure

At the day of sacrifice, animals were deeply anesthetized with isoflurane and sacrificed by decapitation. Trunk blood was collected in labeled 1.5 ml EDTA-coated microcentrifuge tubes (Sarstedt, Germany) and kept on ice until centrifugation. After centrifugation (4°C, 8000rpm for 1 min) the plasma was removed and transferred to new, labeled tubes and stored at −20°C until hormone quantification. For mRNA analysis, brains were removed, snap-frozen in isopentane at −40°C and stored at −80°C for ISH. For protein analysis, brains were removed and placed inside a brain matrix with the hypothalamus facing upwards (spacing 1 mm, World Precision Instruments, Berlin, Germany). Starting from the middle of the chiasma opticum, a 1 mm thick brain slice was removed. The PVN was further isolated by cutting the slice on both sides of the PVN along the formices (parallel to the 3^rd^ ventricle) as a landmark and a horizontal cut between the reuniens and the thalamic nucleus. Finally, the slice containing the 3^rd^ ventricle was bisected, and the distal part discarded. The remaining part (containing the PVN) was immediately shock frozen and stored at −80°C until protein analysis^42^. The adrenals and thymus glands were dissected from fat and weighed.

### Hormone assessment

CORT and ACTH concentrations were determined by radioimmunoassay using a corticosterone double antibody ^125^I RIA kit (sensitivity: 12.5 ng/ml, MP Biomedicals Inc) and adrenocorticotropic double antibody hormone ^125^I RIA kit (sensitivity: 10 pg/ml, MP Biomedicals Inc) and were used following the manufacturers’ instructions. Radioactivity of the pellet was measured with a gamma counter (Packard Cobra II Auto Gamma; Perkin-Elmer). Final CORT and ACTH levels were derived from the standard curve.

### *In-situ* hybridization

ISH was used to analyze mRNA expression of the major stress markers, *Fkbp5*, *Gr*, *Crh*, and *Avp*. Therefore, frozen brains were sectioned at −20 °C in a cryostat microtome at 20 μm, thaw mounted on Super Frost Plus slides, dried and stored at −80 °C. The ISH using ^35^S UTP labeled ribonucleotide probes was performed as described previously ^21,29^. All primer details are available upon request. Briefly, sections were fixed in 4% paraformaldehyde and acetylated in 0.25% acetic anhydride in 0.1 M triethanolamine/HCl. Subsequently, brain sections were dehydrated in increasing concentrations of ethanol. The antisense cRNA probes were transcribed from a linearized plasmid. Tissue sections were saturated with 100 μl of hybridization buffer containing approximately 1.5 × 10^6^ cpm ^35^S labeled riboprobe. Brain sections were coverslipped and incubated overnight at 55°C. On the next day, the sections were rinsed in 2 × SSC (standard saline citrate), treated with RNAse A (20 mg/l). After several washing steps with SSC solutions at room temperature, the sections were washed in 0.1 × SSC for 1 h at 65°C and dehydrated through increasing concentrations of ethanol. Finally, the slides were air-dried and exposed to Kodak Biomax MR films (Eastman Kodak Co., Rochester, NY) and developed. Autoradiographs were digitized, and expression was determined by optical densitometry utilizing the freely available NIH ImageJ software. The mean of four measurements of two different brain slices was calculated for each animal. The data were analyzed blindly, always subtracting the background signal of a nearby structure not expressing the gene of interest from the measurements. For *Fkbp5*, slides were dipped in Kodak NTB2 emulsion (Eastman Kodak Co., Rochester, NY) and exposed at 4 °C for three weeks for better visibility of the viral spread and genetic deletion. Slides were developed and examined with a light microscope with darkfield condensers to show mRNA expression.

### RNAscope analysis and cell counting

For the RNAscope experiments, C57BL/6J male mice were obtained from The Jackson Laboratory (Bar Harbour, ME, USA). All procedures conformed to National Institutes of Health guidelines and were approved by McLean Hospital Institutional Animal Care and Use Committee. Mice were housed in a temperature-controlled colony in the animal facilities of McLean Hospital in Belmont, MA, USA. All mice were group-housed and maintained on a 12:12 h light/dark cycle (lights on at 07:00 am). Food and water were available ad libitum unless specified otherwise. Mice were 12 weeks at the time of tissue collection. Animals were allowed to acclimate to the room for 1 week before the beginning of the experiment. During the experiment mice were either left undisturbed (ctrl) or subjected to 14 hrs (overnight) of food deprivation (FD) prior to sacrifice. During the stress procedure, animals were kept in their home cages and had free access to tap water. All mice were sacrificed by decapitation in the morning (08:00 to 08:30 am) following quick anesthesia by isoflurane. Brains were removed, snap-frozen in isopentane at −40°C, and stored at −80°C. Frozen brains were sectioned in the coronal plane at −20°C in a cryostat microtome at 18 μm, mounted on Super Frost Plus slides, and stored at −80°C. The RNA Scope Fluorescent Multiplex Reagent kit (cat. no. 320850, Advanced Cell Diagnostics, Newark, CA, USA) was used for mRNA staining. Probes used for staining were; mm-Avp-C3, mm-Crh-C3, mm-Fkbp5-C2, mm-Oxt-C3, mm-Sst-C3, and mm-Trh-C3. The staining procedure was performed according to manufacturer’s specifications. Briefly, sections were fixed in 4% paraformaldehyde for 15 min at 4°C. Subsequently, brain sections were dehydrated in increasing concentrations of ethanol. Next, tissue sections were incubated with protease IV for 30 min at room temperature. Probes (probe diluent (cat. no. 300041 used instead of C1-probe), Fkbp5-C2 and one of the above C3-probes) were hybridized for 2 hrs at 40°C followed by 4 hybridization steps of the amplification reagents 1 to 4. Next, sections were counterstained with DAPI, cover-slipped and stored at 4°C until image acquisition. Images of the PVN (left and right side) were acquired by an experimenter blinded to the condition of the animals. Sixteen-bit images of each section were acquired on a Leica SP8 confocal microscope using a 40× objective (n=3 animals per marker and condition). For every individual marker, all images were acquired using identical settings for laser power, detector gain, and amplifier offset. Images of both sides were acquired as a z-stack of 3 steps of 1.0 μm each. *Fkbp5* mRNA expression and co-expression was analyzed using ImageJ with the experimenter blinded to condition of the animals. *Fkbp5* mRNA was counted manually and each cell containing 1 mRNA dot was counted as positive.

### Single-cell sequencing

#### Tissue dissociation

Mice were anesthetized lethally using isoflurane and perfused with cold PBS. Brains were quickly dissected, transferred to ice-cold oxygenated artificial cerebral spinal fluid (aCSF), and kept in the same solution during dissection. Sectioning was performed using a 0.5 mm stainless steel adult mouse brain matrice (Kent Scientific) and a Personna Double Edge Prep Razor Blade. A slide (approximately −0.58 mm Bregma to −1.22 mm Bregma) was obtained from each brain and the extended PVN was manually dissected under the microscope. The PVN from five different mice was pooled and dissociated using the Papain dissociation system (Worthington) following the manufacturer’s instructions. All solutions were oxygenated with a mixture of 5% CO_2_ in O_2_. After this, the cell suspension was filtered with 30 μm filter (Partec) and kept in cold and oxygenated aCSF.

#### Cell capture, library preparation, and high-throughput sequencing

Cell suspensions of PVN with approximately 1.000.000 cells/μL were used. Cells were loaded onto two lanes of a 10× Genomics Chromium chip per factory recommendations. Reverse transcription and library preparation was performed using the 10× Genomics Single Cell v2.0 kit following the 10× Genomics protocol. The library molar concentration and fragment length was quantified by qPCR using KAPA Library Quant (Kapa Biosystems) and Bioanalyzer (Agilent high sensitivity DNA kit), respectively. The library was sequenced on a single lane of an Illumina HiSeq4000 system generating 100bp paired-end reads at a depth of ~340 million reads per sample.

#### Quality control and identification of cell clusters

Pre-processing of the data was done using the 10× Genomics Cell Ranger software version 2.1.1 in default mode. The 10X Genomics supplied reference data for the mm10 assembly and corresponding gene annotation was used for alignment and quantification. All further analysis was performed using SCANPY version 1.3.7 ^43^. A total of 5.113 cells were included after filtering gene counts (<750 and >6.000), UMI counts (>25.000) and fraction of mitochondrial counts (>0.2). Combat^44^ was used to remove chromium channel as batch effect from normalized data. The 4.000 most variable genes were subsequently used as input for Louvain cluster detection. Cell types were determined using a combination of marker genes identified from the literature and gene ontology for cell types using the web-based tool: mousebrain.org (http://mousebrain.org/genesearch.html).

#### Western blot analysis

Protein extracts were obtained by lysing cells (in RIPA buffer (150 mM NaCl, 1% IGEPAL CA-630, 0.5% Sodium deoxycholate, 0.1% SDS 50mM Tris (pH8.0)) freshly supplemented with protease inhibitor (Merck Millipore, Darmstadt, Germany), benzonase (Merck Millipore), 5 mM DTT (Sigma Aldrich, Munich, Germany), and phosphatase inhibitor (Roche, Penzberg, Germany) cocktail. Proteins were separated by SDS-PAGE and electro-transferred onto nitrocellulose membranes. Blots were placed in Tris-buffered saline, supplemented with 0.05% Tween (Sigma Aldrich) and 5% non-fat milk for 1 h at room temperature and then incubated with primary antibody (diluted in TBS/0.05% Tween) overnight at 4°C. The following primary antibodies were used: Actin (1:5000, Santa Cruz Biotechnologies, sc-1616), GR (1:1000, Cell Signaling Technology, #3660), p-GR Ser211 (1:500, Sigma, SAB4503820), p-GR Ser226 (1:1000, Sigma, SAB4503874), p-GR 203 (1:500, Sigma, SAB4504585), FKBP51 (1:1000, Bethyl, A301-430A).

Subsequently, blots were washed and probed with the respective horseradish peroxidase or fluorophore-conjugated secondary antibody for 1 h at room temperature. The immuno-reactive bands were visualized either using ECL detection reagent (Millipore, Billerica, MA, USA) or directly by excitation of the respective fluorophore. Determination of the band intensities were performed with BioRad, ChemiDoc MP. For quantification of phosphorylated GR, the intensity of phosphor-GR was always referred to the signal intensity of the corresponding total GR.

#### Chromatin preparation for chromatin immunoprecipitation (ChIP) analysis

The GR ChIP was performed as previously described^26^. We added 1 mM AEBSF or 0.1 mM PMSF, 5mM Na^+^-Butyrate (NaBut), and PhosSTOP phosphatase inhibitor cocktail tablets (1 per 10 ml; Roche, Burgess Hill, UK) to all solutions unless otherwise stated. Briefly, hypothalamus tissues from four mice were cross-linked for 10 min in 1% formaldehyde in PBS. Crosslinking was terminated by adding glycine (5 min, final concentration 200 μM) and centrifuged (5 min, 6000 g, 4°C). Pellets were washed three times with ice-cold PBS. Next, the pellets were re-suspended in ice-cold Lysis Buffer [50 mM Tris-HCl pH 8, 150 mM NaCl, 5mM EDTA pH 8.0, 0.5% v/v Igepal, 0.5% Na-deoxycholate, 1% SDS, 5mM NaBut, 2 mM AEBSF, 1mM Na3VO4, Complete ultra EDTA-free protease inhibitor tablets and PhosSTOP phosphatase inhibitor cocktail tablet (both 1 per 10 ml, Roche, Burgess Hill, UK)] and rotated for 15 min at 4°C. Samples were aliquoted, sonicated (high power; 2 × 10 cycles; 30 s ON, 60 s OFF) using a water-cooled (4°C) Bioruptor Pico (Diagenode, Liège, Belgium) and centrifuged (10 min, 20,000 g, 4°C). Supernatants (containing the sheared chromatin) were recombined and re-aliquoted into fresh tubes for subsequent ChIP analysis and for assessment of Input DNA (i.e., the starting material). Chromatin was sonicated to a length of approximately 500 base pairs.

#### For ChIP analysis

aliquots of chromatin were diluted 10-times in ice-cold dilution buffer [50 mM Tris-HCl pH 8.0, 150 mM NaCl, 5 mM EDTA pH 8.0, 1% v/v Triton, 0.1% Na-deoxycholate 5mM NaBut, 1 mM AEBSF, Complete Ultra EDTA-free protease inhibitor tablets and PhosSTOP phosphatase inhibitor cocktail tablet (both 1 per 10 ml, Roche)]. 10 μl of GR antibody (ProteinTech, USA) was added to each sample and tubes were rotated overnight at 4°C. Protein A-coated Dynabeads® (Life Technologies) were washed once in ice-cold 0.5% BSA/PBS before blocking overnight at 4°C. Pre-blocked beads were washed once in ice-cold dilution buffer, re-suspended in the antibody:chromatin mix, and allowed to incubate for 3 h at 4°C to allow binding of beads to antibody:chromatin complexes. After 3 h, the samples were placed in a magnetic stand to allow the beads (with the Bound fraction bound) to separate from the liquid ‘Unbound’ fraction. Beads carrying the Bound chromatin were washed 3 times with ice-cold RIPA buffer [10 mM Tris-HCl pH 7.5, 1 mM EDTA pH 7.5, 0.1% SDS, 0.5 mM EGTA, 1% Triton, 0.1% Na-Deoxycholate, 140 mM NaCl + inhibitors] and washed twice with ice-cold Tris-EDTA buffer. Bound DNA was eluted in two steps at room temperature; first with 200 μl Elution buffer 1 (10 mM Tris-HCl pH 7.4, 50 mM NaCl, 1.5% SDS) and second with 100 μl Elution buffer 2 (10 mM Tris-HCl pH 7.4, 50 mM NaCl, 0.5% SDS). Crosslinks were reversed by addition of NaCl (final concentration 200 mM) and overnight incubation at 65°C. The next day, samples were incubated first with RNase A (60 μg/ml, 37°C, 1 h), followed by incubation with proteinase K (250 μg/ml, 37°C, 3.5 h). DNA was purified using a QIAquick PCR purification kit (Qiagen) as per manufacturer’s instructions. Input samples were incubated overnight at 65°C with 200 mM NaCl to reverse crosslinks, incubated with RNase A and proteinase K (overnight), and DNA was purified using a Qiagen PCR purification kit. All samples (Bounds and Inputs) were diluted to a standardized concentration with nuclease free water and analyzed by qPCR as described below using primers/probes (Forward: 5’– TGTCAATGGACAAGTCATAAGAAACC; Reverse: 5’ – GAATCTCACATCCAATTATATCAACAGAT; Probe: 5’ – TTCCATTTTCGGGCTCGTTGACGTC. Binding of GR was expressed as a percentage of input DNA, i.e., % Input, which is a measure of the enrichment of steroid receptor bound to specific genomic sequences.

#### qPCR analysis

Mastermix for qPCR was prepared containing 900 nM forward and reverse primers, 200 nM probe, 1× TaqMan fast mastermix (Life Technologies, Paisley, UK) and made up to volume with nuclease-free water. Primers and dual-labeled probe with 6-FAM as the fluorescent dye and TAMRA as the quencher were designed using Primer Express software (Version 3.0.1, Life Technologies). Standard curves were performed for each primer pair and the qPCR efficiency was calculated using the equation: E = ((10-1/slope) − 1) × 100 (where E is qPCR efficiency and the slope is the gradient of the standard curve). Only primer pairs with efficiencies greater than 90% were used. Quantitative PCR was performed using a StepOne Plus machine (Life Technologies, Paisley, UK). Taq enzymes were activated at 95°C for 20 s, then 40 cycles of 95°C (1 s) to 60°C (20 s) were performed to amplify samples.

#### Statistical analysis

The data presented are shown as means ±SEM and samples sizes are indicated in the figure legends. All data were analyzed by the commercially available software SPSS 17.0 and GraphPad 8.0. When two groups were compared, the unpaired student’s *t*-test was applied. If data were not normally distributed the non-parametric Mann-Whitney test (MW-test) was used. For four group comparisons, two-way analysis of variance (ANOVA) was performed, followed by posthoc test, as appropriate. P values of less than 0.05 were considered statistically significant.

## Acknowledgments

The authors thank Claudia Kühne, Mira Jakovcevski, Daniela Harbich and Bianca Schmid (Max Planck Institute of Psychiatry, Munich, Germany) for their excellent technical assistant and support. We thank Stefanie Unkmeir, Sabrina Bauer and the scientific core unit *Genetically Engineered Mouse Models* for genotyping support. This work was supported by the “OptiMD” grant of the Federal Ministry of Education and Research (01EE1401D; M.V.S.), the BioM M4 award “PROCERA” of the Bavarian State Ministry (M.V.S.) and by a NARSAD Young Investigator Grant from the Brain & Behavior Research Foundation (J.H.).

## Author contributions

A.S.H, and M.V.S.: Conceived the project and designed the experiments. J.M.D.: Provided scientific expertise for establishing *Fkbp5* mouse lines. A.S.H and L.M.B. managed the mouse lines and genotyping. A.S.H., M.L.P., L.R., and L.M.B. performed animal experiments and surgeries. R.S.: Performed CORT and ACTH hormone assays and analysis. J.P.L., S.R., and E.B.: Performed single cell sequencing experiments and analysis. J.M.H.M.R, and H.M.G.: Performed and analyzed GR-CHIP experiments. A.S.H, J.H., and C.E.: Performed and designed RNAscope experiments and manual counting of cells. N.C.G, and K.H.: Performed protein analysis. K.J.R. and A.C.: Supervised and revised the manuscript. A.S.H.: Wrote the initial version of the manuscript. M.V.S.: Supervised the research and all authors revised the manuscript.

## Competing interests

The authors declare no competing interests.

## Supplementary Information

**Suppl. Figure 1:**
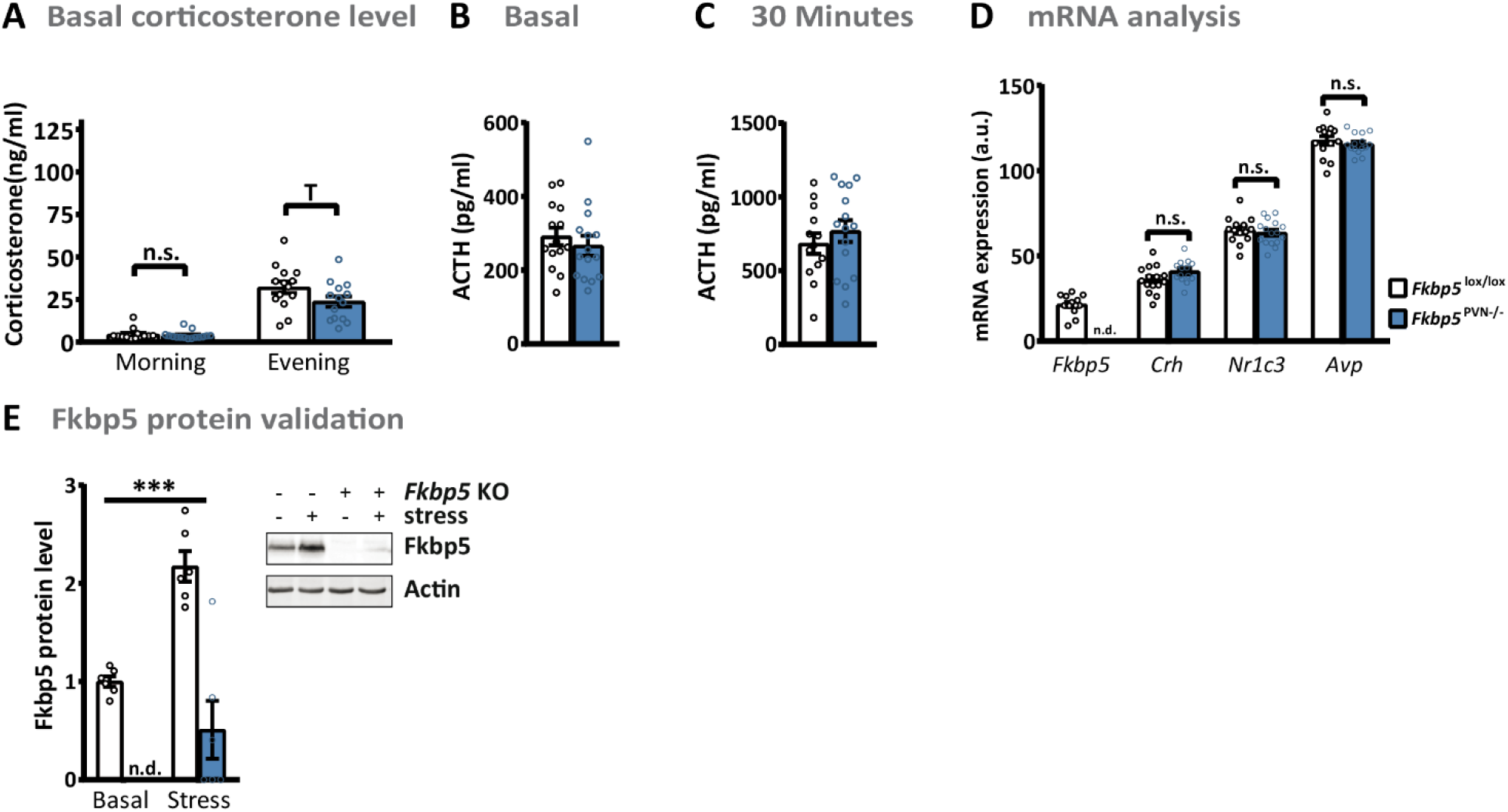
Corticosterone and ACTH levels of *Fkbp5*^PVN−/−^ mice. *Fkbp5* deletion in the PVN has no effect basal CORT levels (*Fkbp5*^PVN−/−^ n = 16; *Fkbp5*^lox/lox^ n =15) **(A).** ACTH levels under basal (*Fkbp5*^PVN−/−^ n = 16; *Fkbp5*^lox/lox^ n =15) **(B)** and 30 minutes after stress onset were unaltered (*Fkbp5*^PVN−/−^ n = 16; *Fkbp5*^lox/lox^ n =15) **(C)** . **(D)** mRNA changes of stress responsive genes within the PVN (*Fkbp5*^PVN−/−^ n = 16; *Fkbp5*^lox/lox^ n =15). **(E)** Fkbp5 protein levels were not detectable under basal conditions. All data are presented as mean ± SEM and were analyzed with a student’s t-test. n.d. = not detectable; n.s. = not significant; T = 0.05 < p < 0.1.

**Suppl. Figure 2:**
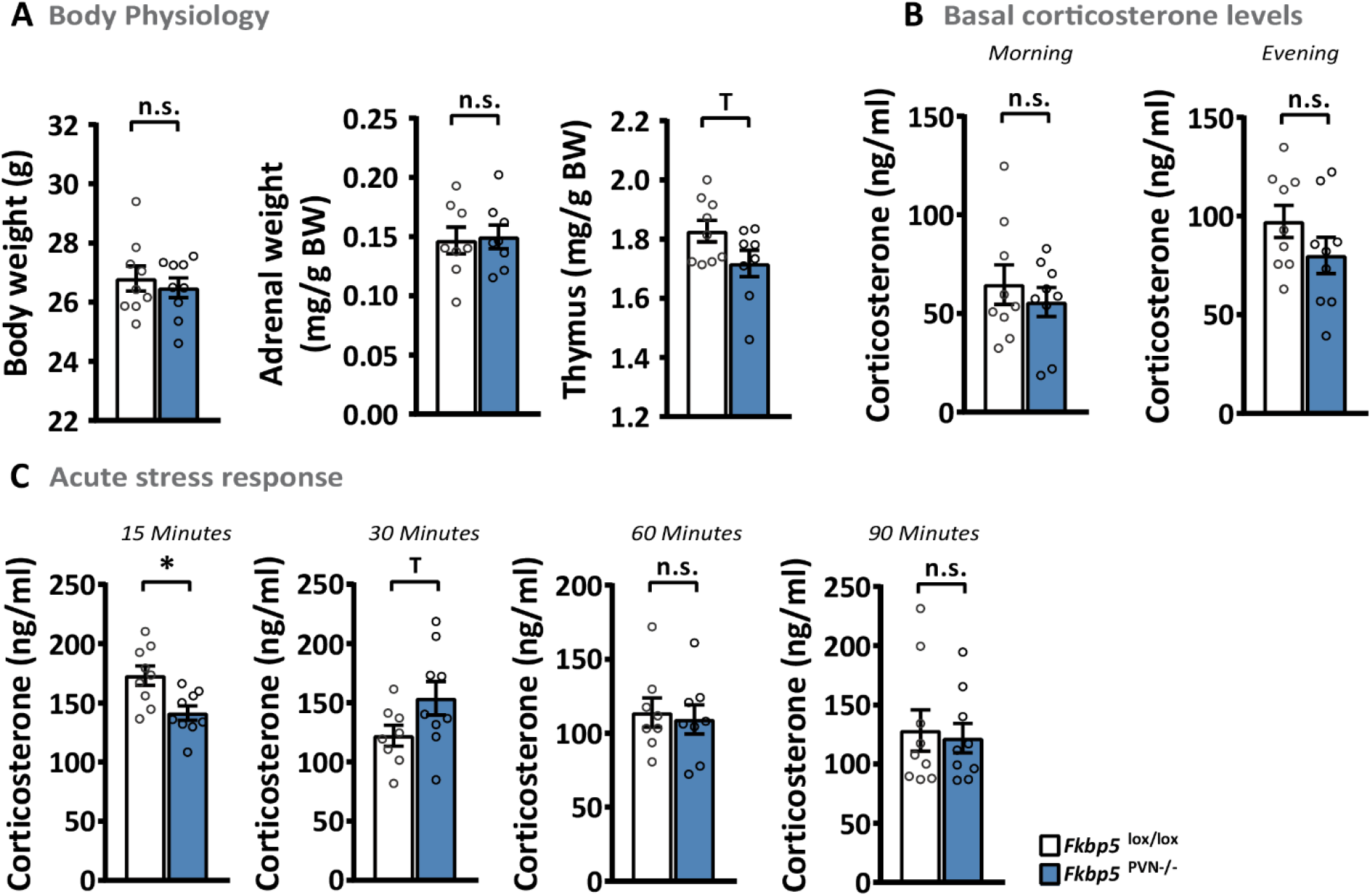
Young mice with *Fkbp5* deletion in the PVN. **(A)** Animals with an age of 10 weeks had no alterations in body physiology. **(B)** Morning and evening corticosterone levels were unchanged. **(C)** *Fkbp5*^PVN−/−^ had significant lower corticosterone levels 15 minutes after stress onset. (Group size for A-C: *Fkbp5*^PVN−/−^ n = 9; *Fkbp5*^lox/lox^ n =9). All data are presented as mean ± SEM and were analyzed with a student’s t-test. n.s. = not significant; T = 0.05 < p < 0.1; * = p < 0.05.

**Suppl. Figure 3:**
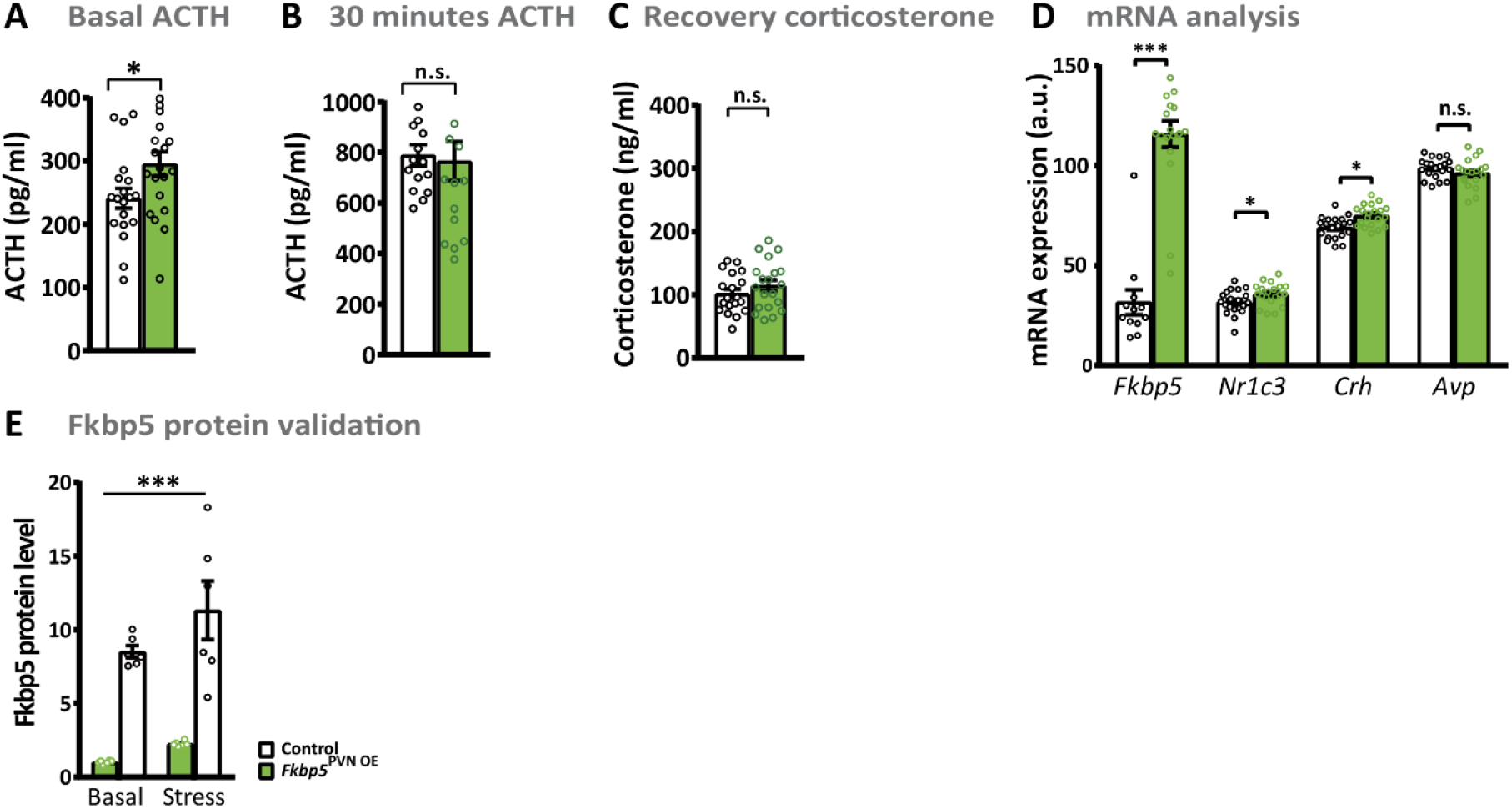
Overexpression of *Fkbp5* in the PVN affected ACTH and mRNA levels. **(A-B)** ACTH levels were significantly higher in *Fkbp5*^PVN OE^ mice under basal (n = 20 vs. 20) and unchanged 30 minutes after stress (n = 12 vs. 12= onset compared to their controls**(C)**. We did not detect any differences in corticosterone levels 90 minutes after stress onset (n = 20 vs. 20). **(D)** *Fkbp5* overexpression resulted in significant increase in mRNA levels of *Fkbp5*, *N1c3* and *Crh*. *Avp* levels stayed unchanged (n = 20 vs. 20). **(E)** Viral overexpression resulted in a 4-fold Fkbp5 protein upregulation in the PVN (n = 20 vs. 20). All data are presented as mean ± SEM and were analyzed with a student’s t-test (A-D) or with a two way ANOVA (E). n.s. = not significant; T = 0.05 < p < 0.1. * = p < 0.05, *** = p < 0.001.

**Suppl. Figure 4:**
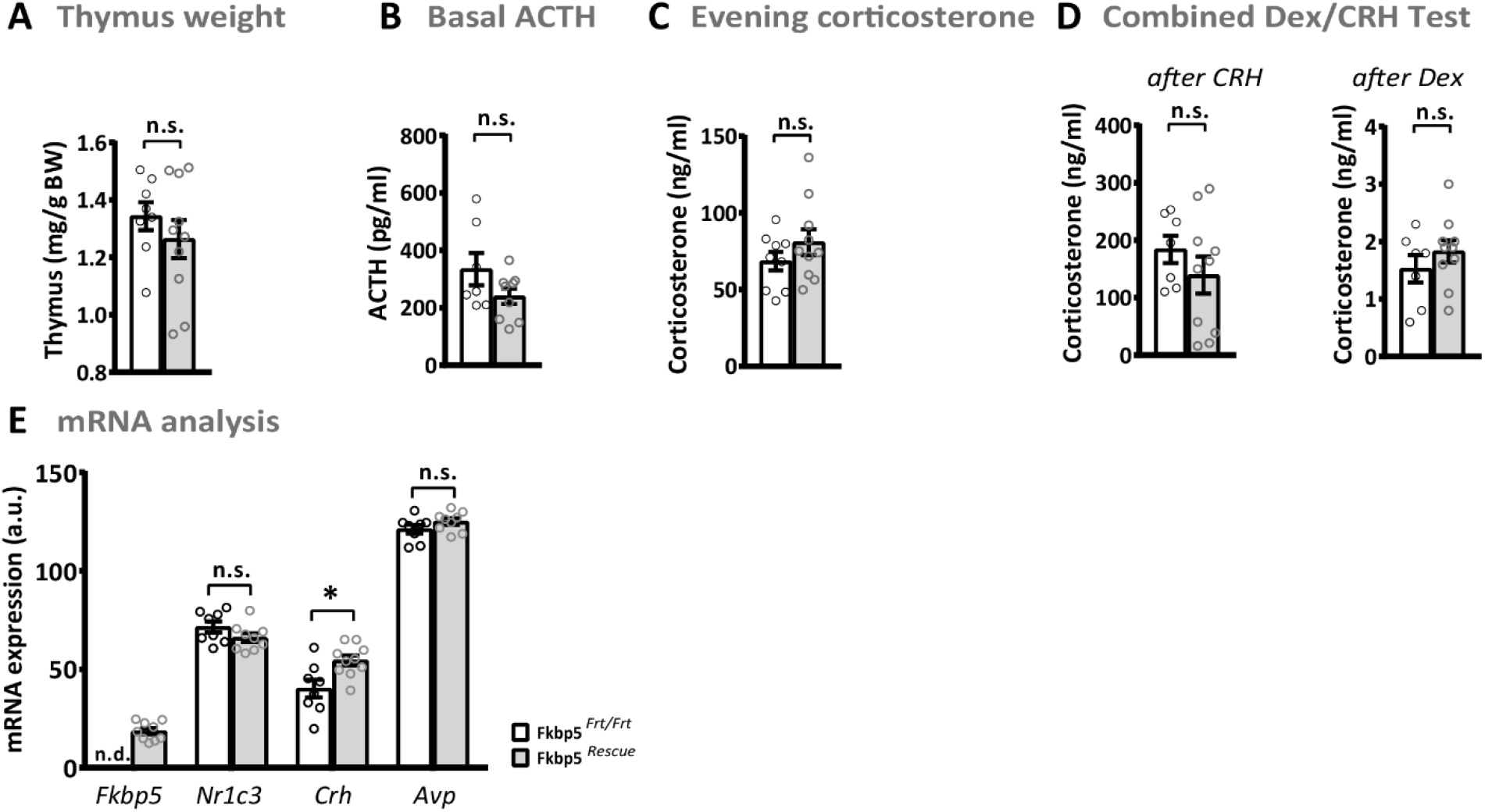
Corticosterone and ACTH levels of *Fkbp5*^Rescue^ mice. *Fkbp5* reinstatement had no effect on thymus weights **(A)**. Basal ACTH and evening corticosterone levels were unaltered **(B & C)**. **(D)**Rescue of endogenous *Fkbp5* in global knock-out animals had no significant effect on the Dex/CRH test and on stress responsive mRNA levels in the brain. **(E)** mRNA levels in *Fkbp5*^Rescue^ animals compared to global knock-out littermates. Group sizes for A-E: *Fkbp5*^Rescue^ n = 10; *Fkbp5*^Frt/Frt^ n =9) All data are presented as mean ± SEM and were analyzed with a student’s t-test. n.s. = not significant.

**Suppl. Figure 5:**
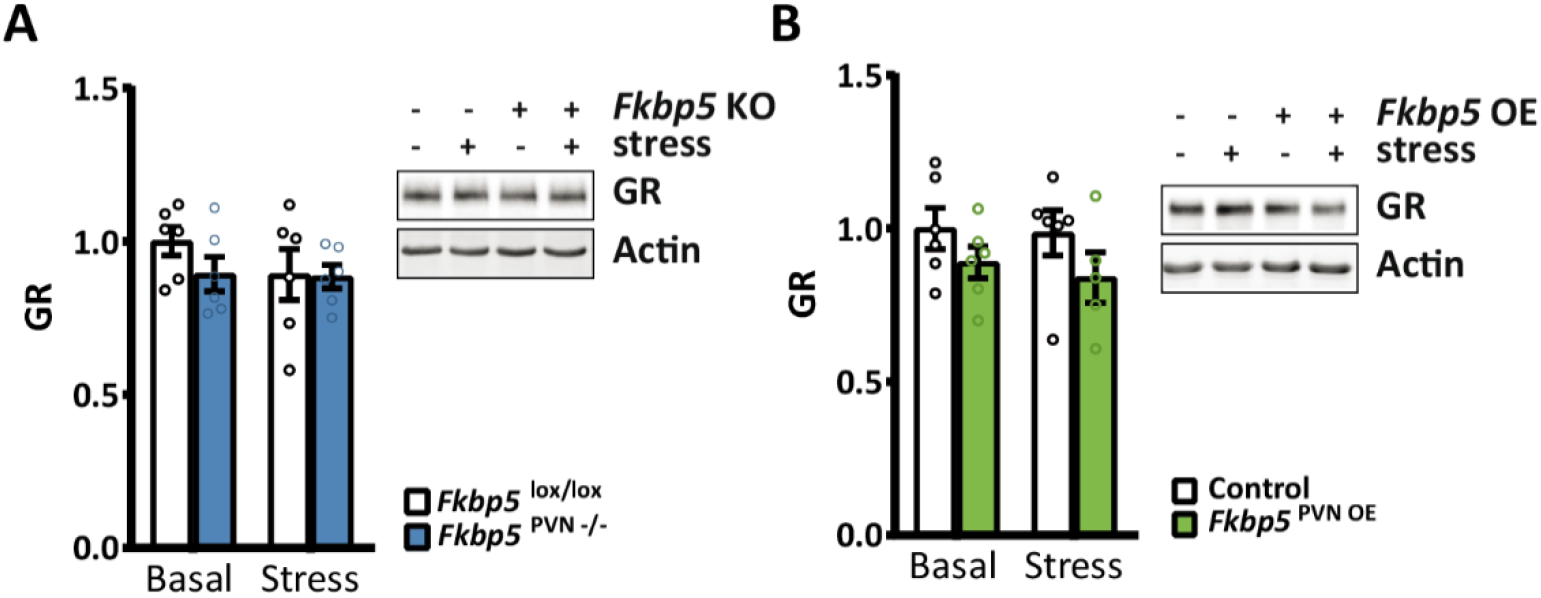
PVN protein level of Fkbp5 and GR under basal and stressed conditions. **(A)** GR protein levels were unchanged in *Fkbp5*^PVN−/−^ animals (n = 6) and **(B)** in *Fkbp5*^PVN OE^ animals (n = 6) under basal and stressed conditions. All data were analyzed with a two-way ANOVA.

**Suppl. Figure 6:**
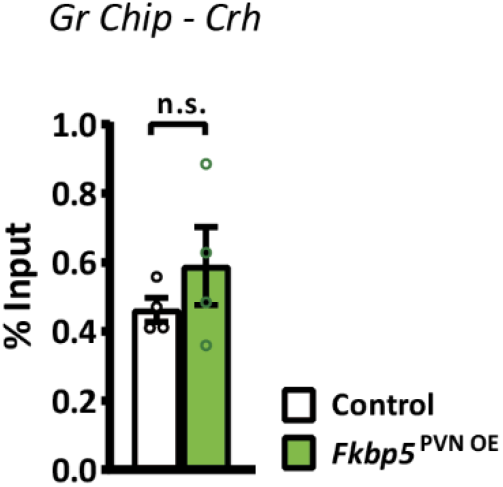
GR to GRE binding within the *Crh* gene after stress. Mice were sacrificed 30 minutes after stress onset. Every n consists of a pool of 4 individual hypothalami. All data are indicated as mean ± SEM and were analyzed with a student’s t-test; (n = 4 vs. 4); n.s. = not significant.

**Suppl. Figure 7:**
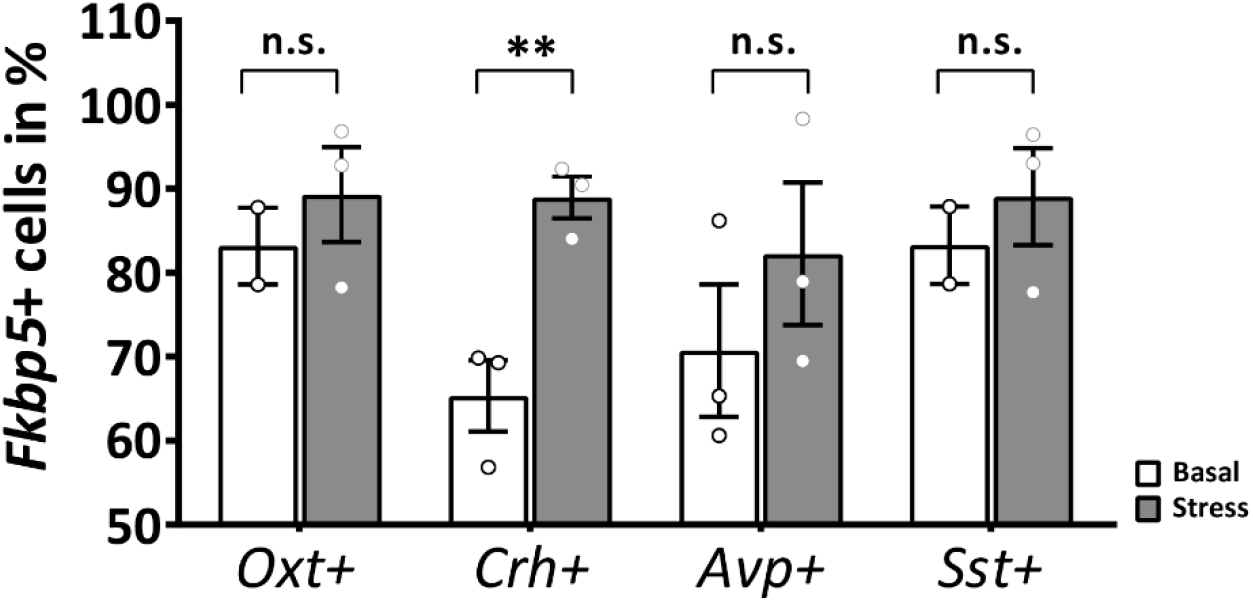
Quantification of *Fkbp5* co-localization in stress response markers under basal and stress conditions. Under basal conditions *Fkbp5* mRNA signal was detected in 83% of *Oxt*+ neurons, 65% Of *Crh*+ neurons, 68% in *Avp*+ and 84% of *Sst*+ neurons. We monitored an increased in *Fkbp5* mRNA expression in all neuronal populations. Interestingly, only *Crh*+ neurons showed a significant increase of *Fkbp5* mRNA following stress. All data are presented as mean ± SEM and were analyzed with a student’s t-test. ** = p < 0.01.

